# 16S rRNA sequence captures microbial functional potential

**DOI:** 10.64898/2026.04.16.718937

**Authors:** Jia Liu, M. Clara De Paolis Kaluza, Yana Bromberg

## Abstract

16S rRNA amplicon sequencing is widely used for microbiome profiling, but most methods rely on reference databases of characterized organisms, limiting its accuracy in function prediction for underrepresented environments. We discovered that 16S rRNA k-mer composition carries substantial functional signal: (i) whole-genome k-mer profiles predict genome-encoded functions, and (ii) 16S rRNA k-mer profiles reflect their source genome’s composition. Building on these relationships, we developed embeRNA, a neural network framework that predicts functions directly from 16S rRNA k-mer embeddings without requiring taxonomy assignment or phylogenetic placement. embeRNA outputs per-function probability scores, enabling users to tune decision thresholds to balance precision and recall or account for community novelty. In a stringent “novel microbes” benchmark - where all test sequences shared <97% identity with training data - embeRNA outperformed reference-based methods, particularly for hard-to-label functions. Applied to soil metagenomes with paired 16S and whole metagenome shotgun sequencing (WMS) data, embeRNA recovered most WMS-inferred functions and produced abundance profiles strongly correlated with WMS results, attaining better performance than a reference-based approach. Our findings demonstrate that 16S rRNA directly captures functional potential, and 16S amplicon sequencing data can complement WMS-based inference to broaden functional characterization of microbiomes, especially in understudied environments.

## Introduction

Microbial communities inhabit every ecosystem and host-associated environment^1–3^, where they collectively drive core biochemical reactions and shape ecological processes^4–6^. Advances in whole metagenome shotgun (WMS) sequencing and targeted 16S rRNA amplicon sequencing have enabled broad surveys of community compositions and, increasingly, functional potentials.

A wide range of computational pipelines (at least 49 by our count) are routinely used to translate sequencing data into taxonomic profiles and/or functional potentials^7–55^ (**Figure 1**). Whole metagenome shotgun (WMS) sequencing -inferred functions are generally treated as being high-confidence/precision. However, WMS sequencing depth limits coverage of individual, especially rare, genes and organisms, making it easy to miss low-abundance or infrequent functionality (low recall) and limiting the resolution of abundance estimates^56–58^.

**Figure 1.**
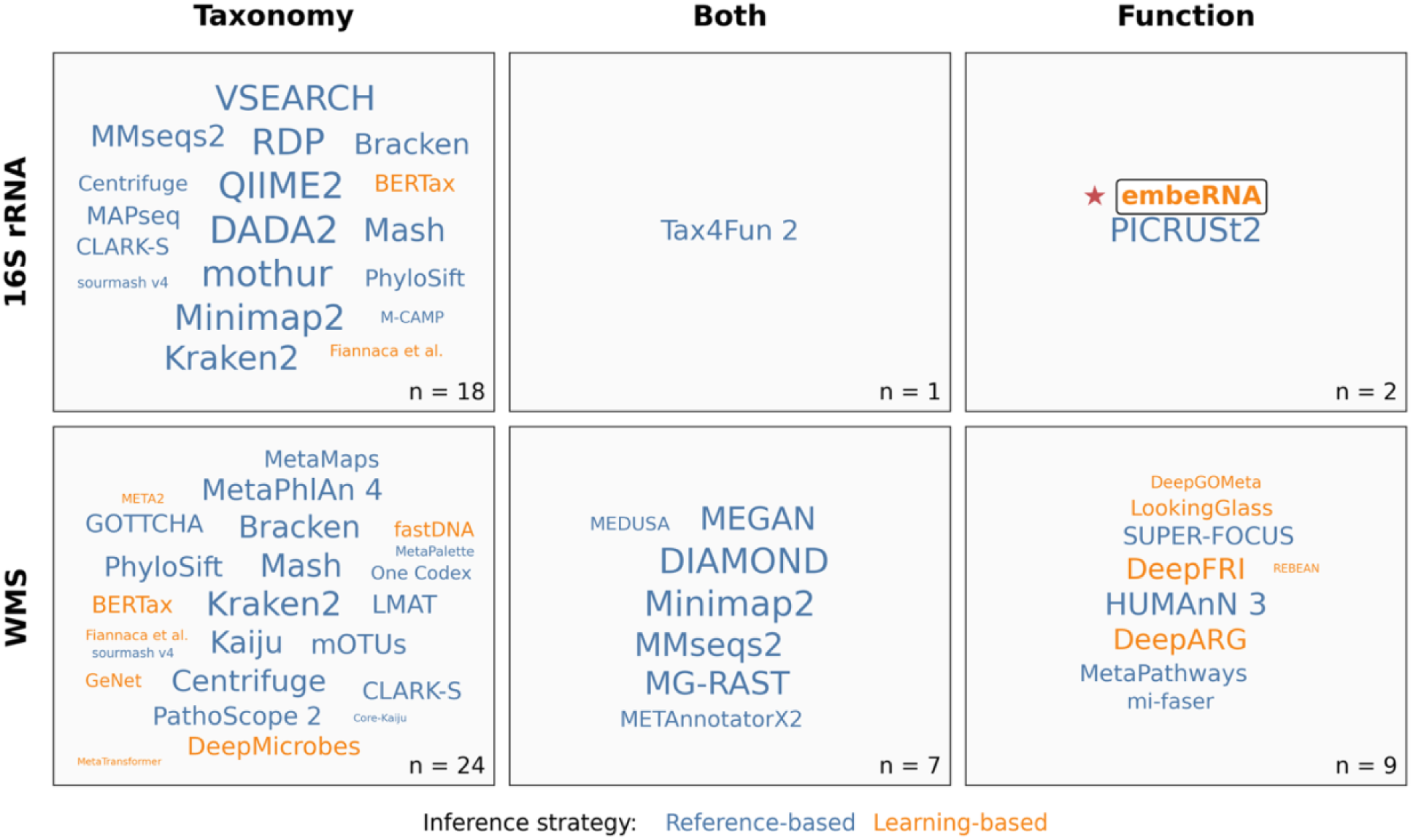
Microbiome analysis tools vary. 49 routinely used microbiome analysis tools are organized by input sequence type (16S rRNA or whole metagenome shotgun sequencing, WMS) and primary prediction target (taxonomy, both taxonomy and function, or function). For existing tools, name size is proportional to log2-transformed Google Scholar citation count; embeRNA is highlighted as the method introduced in this study and is not citation-scaled. Color indicates inference strategy (blue: reference-based tools or tools commonly embedded in workflow requiring reference databases; orange: learning-based tools). Reference-based methods infer outputs by anchoring input data to known references, whereas learning-based methods infer outputs from patterns learned during training without explicit reference matching at inference time.Because some tools, such as Diamond or MMseqs2, are used broadly beyond microbiome research, citation count/name size should be interpreted as a rough indicator of overall adoption rather than microbiome-specific usage. Alt text: Grid of six panels organizing 49 microbiome analysis tools by input data type and prediction target. Tool names are scaled by citation count, coloured blue for reference-based and orange for learning-based methods. embeRNA is highlighted with a star as a learning-based tool in the 16S rRNA × Function cell, which contains only two tools, the other being reference-based.

16S rRNA amplicon sequencing is a cost-effective approach for ensuring both high precision and recall in profiling microbiome taxonomic composition, typically at the genus level. Existing annotation methods link 16S rRNA sequences to the corresponding microbial isolates via aligning input sequences^18^ or corresponding k-mer vectors^39, 59^ to reference trees or databases^59–61^. Once microbiome taxonomic/phylogenetic profiles are established, corresponding functional profiles can be inferred by combining functions of member taxa, e.g. as is done by PICRUSt / PICRUSt2 or Tax4Fun^40, 62, 63^. While this approach is effective for well-characterized environments such as the human gut^64, 65^, it is constrained by reference data availability, yielding biased or incomplete function predictions for poorly characterized microbiomes.

To summarize, most extant approaches focus on resolving taxonomy, whether from 16S or WMS data (**Figure 1**). Many methods treat taxonomic composition as an end goal, answering the question “Who is there?” Some determine the taxonomic composition as the first step in functional inference, asking “What are they doing?” Notably, most of these tools rely on reference-based inference, or are commonly embedded in workflows that depend on reference databases, whereas learning-based approaches infer outputs from learned sequence patterns without explicit reference matching during inference. In contrast, some methods profile functions from WMS data directly, where both reference-based and learning-based approaches are available. Overall, however, accurately and comprehensively characterizing microbiome functional potential remains a challenge. Here we ask, for the first time, can microbiome functions be inferred from 16S rRNA sequence directly, i.e. without seeking a reference?

### Sequence is key to understanding function

Short k-mer composition vectors provide compact, fixed-size representations of DNA sequences of any length, capturing reproducible genomic signatures such as GC content and oligonucleotide usage biases^66–69^. Because the 16S rRNA gene is embedded within the genome, its k-mer patterns may reflect aspects of genome-wide sequence composition, resulting from exposure to the same evolutionary constraints and, possibly, carrying information beyond taxonomy alone. Indeed, prior studies have shown that 16S rRNA k-mer features can predict microbiome sample characteristics and microbial phenotypes, e.g. habitat or host-associated traits^70^, suggesting that marker-gene k-mers encode informative biological signal.

Here, we show that (i) whole genome k-mer representations are predictive of microbial functions and that (ii) 16S rRNA k-mer representations reflect corresponding whole genome k-mer representations. Together, these observations support a direct 16S rRNA-to-function link.

To make further use of this link, we built embeRNA (embedding RNA), a neural network-based method that predicts functional profiles directly from 16S rRNA k-mer embeddings without explicit phylogenetic placement, or taxonomic assignments.

## Materials and Methods

### Datasets description

This work used four datasets as described below (***Balanced, embeRNA, Novel Microbes,*** and ***Microbiome sets***).

1. ***Balanced set:*** To assess the 16S rRNA k-mers to genome k-mers to function relationships, we assembled a phylogenetically diverse set of bacterial genomes. Our group previously constructed the Fusion database^71^, comprising 433,891 function clusters derived from 8,906 complete bacterial genome assemblies from NCBI RefSeq database. From this resource, a species-balanced subset of 1,502 assemblies was created to capture ≥90% of the genetic diversity present across the full Fusion genome collection^71^. Here, we use this subset, further filtering it to retain assemblies with (i) at least one full-length 16S rRNA gene and (ii) an annotated species-level taxonomy label. This *Balanced set* comprises 1,369 organisms/assemblies spanning 1,349 unique species and containing 4,744 full-length 16S rRNA sequences (median 3 copies per organism; Interquartile range / IQR 2-4). From this resource, we constructed two datasets: (i) 16S rRNA/genome pairs, where each of the 16S rRNA sequences and its source genome were independently represented as concatenated 1, 2, 3, 4 and 5-mer (1to5-mer) vectors; and (ii) genome/function pairs, where each of the 1,369 organisms’ 1to5-mer genome vector was paired with its Enzyme Commission (EC) number^72^ three-digit functional profile. Across all 1369 organisms in the *Balanced set*, 208 unique EC three-digit functions were observed. Individual organisms carried a median of 124 [IQR: 106-137] functions, while each function was present in 806 [IQR: 311-1,281] assemblies.
2. ***embeRNA set:*** To develop models that infer microbial functions directly from 16S rRNA k-mer composition, we compiled *embeRNA set*, a dataset of 24,585 unique complete prokaryotic genome assemblies derived from (i) PICRUSt2^40^ (n = 16,136) and (ii) Fusion^71^ (n = 8,449) references. Note that while all genomes from Fusion database are bacteria, 364 of PICRUSt2 genomes are archaea. Assemblies that appeared in both resources (n = 2,620) were removed from the PICRUSt2 set to avoid duplication. All retained assemblies were complete genomes containing at least one 16S rRNA sequence and assigned taxonomic labels (species to phylum). Overall, the *embeRNA set* spans 10,359 unique species across 2,262 genera (**SOM Figure 1**) and contains 56,734 16S rRNA sequences (median 1 [IQR: 1-3] copy per genome). Each of these 16S rRNAs was represented to 1to5-mer, and was paired with the EC (three-digit) function profile of its source genome. Because the original PICRUSt2 EC profiles do not include EC numbers from enzymatic class seven^40^, we re-extracted full-digit EC annotations for all PICRUSt2 reference genomes from the JGI IMG database^73^ using the same procedure employed in PICRUSt2^40^, and subsequently summarized these annotations to EC three-digit profiles. When identical 16S rRNA sequences were associated with different annotated function sets across assemblies, we retained each sequence with its original function profile. Across the 8,449 Fusion assemblies, 209 unique EC three-digit functions were observed, spanning all seven enzymatic classes. The 16,136 PICRUSt2 assemblies contained 260 EC three-digit functions, also spanning all seven classes. We defined functional space of *embeRNA set* as the union of the above two, yielding 261 EC three-digit functions, corresponding to 81% of all existing EC three-digit classes recognized at the time of writing (accessed 01/16/2026)^74^. Within this functional space, a typical genome has a median of 141 [IQR: 122-155] EC three-digit functions, and each function is present in 14,140 [IQR: 2,506-23,385] genomes. For embeRNA model training, we further restricted the target space to EC functions present in at least 1% of all assemblies in *embeRNA set*, resulting in 238 EC three-digit functions used in the embeRNA models. Functions underrepresented in current reference collections fall outside this scope regardless of their true environmental frequency. This threshold is user-definable, allowing users to adjust the precision/recall tradeoff for their specific application.
3. ***Novel Microbes set:*** To evaluate model generalizability, we additionally assembled an evaluation set of 2,533 complete bacterial genomes comprising 5,542 unique full-length 16S rRNA genes deposited into the RefSeq database after (02/28/2018) the construction of both components of the *embeRNA set* (Fusion/PICRUSt2). In this set, each 16S rRNA had <97% sequence identity (typical same species threshold) to any of the sequences in the *embeRNA set* (**SOM Note 1**). Each pair of the 5,542 16S rRNA sequence’ 1to5-mer embedding with the EC three-digit function profile of its source genome was generated and served as one sample pair of input and output. The 2,533 assemblies contain a total of 241 unique EC three-digit functions, spanning all seven enzyme classes.
4. ***Microbiome set:*** To evaluate model performance on real amplicon sequencing data, we used a set of 22 bulk and blueberry rhizosphere soil samples with both 16S rRNA V6-V8 amplicon sequencing and WMS sequencing data from a previous study^75^.

### Data preparation

Here we describe sequence featurization, function profile construction, model training/evaluation, and dataset-specific processing steps required to reproduce each analysis.

#### 1to5-mer embeddings

We represented DNA sequences (full-length 16S rRNA genes, extracted 16S rRNA regions, or whole genomes) using k-mer frequency embeddings constructed by concatenating normalized frequency vectors for k = 1-5. For each k, counts of all possible k-mers were computed across the sequence and normalized by the total number of observed k-mers of that order. The resulting five normalized vectors were concatenated into a 1,364-dimensional feature vector. This representation was used for all sequence-based analyses and models in this study.

#### EC three-digit functional profiles

We represented genome enzymatic potential using EC three-digit presence/absence profiles, derived from four-digit EC annotations for each genome assembly in the *Balanced*, *embeRNA*, and *Novel Microbes* datasets.

Four-digit EC annotations for assemblies in the above-mentioned sets were first collected. From the latest Fusion database, we obtained the four-digit ECs for the Fusion-sourced assemblies in *embeRNA set* and all assemblies in the *Balanced set*^71^. From JGI IMG databases, we collected the latest four-digit EC annotations for the remaining PICRUSt2-sourced assemblies in *embeRNA set*^73^. For assemblies in the *Novel Microbes set*, EC annotations were generated de novo. Protein sequences for each genome were retrieved from NCBI RefSeq and annotated with four-digit EC numbers following the procedure as described in the Fusion study^71^. Protein-level EC annotations were then aggregated at the genome level.

For each genome, four-digit EC numbers were then truncated to the first three digits (e.g., 1.1.1.2 → 1.1.1). An EC three-digit function was considered present if at least one protein in the genome carried the corresponding EC annotation and absent otherwise.

### Predicting genome k-mer composition from 16S rRNA k-mers

To predict a genome’s 1to5-mers from those of its 16S rRNAs, we trained three regression models on the *Balanced set* of assemblies:

1. A dummy mean baseline model predicted the average per whole genome k-mer value across all training-fold samples for all test samples.
2. A ridge regression model was trained to predict genome 1to5-mers, where model input was the principle components (dimension: 256) extracted from the standardized 16S rRNA 1to5-mers. Regularization strength was selected by inner cross-validation on a log-spaced alpha grid (10⁻⁴-10⁴).
3. A random forest model was also trained to map 16S rRNA 1to5mers to that of genome (300 trees; max depth: 30; min samples per leaf: 10).

#### Cross-validation

Because genomes can encode multiple 16S copies, evaluation used grouped 5-fold cross-validation by assembly, assigning all 16S sequences from the same assembly to the same fold. The same folds were reused across models and controls to enable paired comparisons.

#### Permutation controls

To verify that models learned a genuine 16S rRNA to genome mapping, we repeated training under two training-label destructive regimes (within each training fold) while evaluating against the true test labels separately by: (i) randomly shuffling the 16S rRNA - genome pairings (permuted pairs), and (ii) randomly permuting the 1to5-mer feature positions within each genome vector (genome permutation).

#### Evaluation metrics

We evaluated models’ performance on held-out folds using variance-weighted multi-output R², weighting per-feature R² by the variance of that feature across genomes.

### Predicting EC functional profiles from genome k-mers

On the *Balanced set* of assemblies, we trained models to predict presence/absence of each EC three-digit function encoded by a genome using its genome 1to5-mers:

1. We trained a L2-regularized logistic regression (LR) with class-balanced loss. Features were standardized using training-fold mean and standard deviation, with the same transform applied to test folds.
2. An Extra Trees (ET) classifier was trained to predict function presence/absence (300 trees per label).

#### Cross-validation

To avoid evaluating on closely related genomes, we performed grouped 5-fold cross-validation by species, assigning all assemblies of the same species to the same fold. The same folds were reused across models/baselines/controls.

#### Baselines and permutation control

We included baseline models and permutation controls as following: (i) majority baseline which predicted an EC as present for all test assemblies if its training prevalence ≥0.5; (ii) random-prevalence baseline which predicted presence for each EC by sampling Bernoulli(p) with p equal to the label’s training prevalence; and (iii) permutated-label control which independently permuted each EC across training samples before model training (preserving the prevalence while destroying feature label connections), and evaluated the predictions against true test labels.

#### Evaluation metrics

We evaluated the predicted presence/absence for each function against its ground truth on test folds. We define true positives (TP) as EC labels correctly predicted as present, false positives (FP) as EC labels predicted present but absent in the ground truth, and false negatives (FN) as EC labels present in the ground truth but predicted absent. Precision, recall, and F1 for each function were computed as:

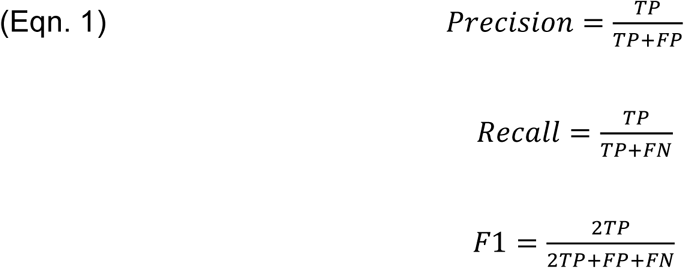

We additionally reported Hamming loss, the fraction of incorrectly predicted labels across all samples and labels within a test fold:

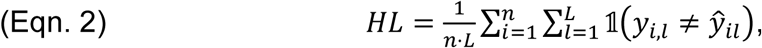

where n is the number of test samples, L is the number of EC labels, 𝑦_𝑖,𝑙_ ∈ {0|1} is the ground-truth presence/absence for label l in sample i, and 𝑦^_𝑖𝑙_ ∈ {0|1} is the corresponding prediction.

### embeRNA: predicting functions from 16S rRNA k-mer

Next, we developed embeRNA framework to inform functions from 16S rRNA k-mer using the *embeRNA set*.

#### 16S rRNA region extraction and embeddings

From each full-length 16S rRNA sequence in the *embeRNA set*, we extracted commonly sequenced regions (V1-V2, V3-V4, V4, V4-V5, V6-V8, V7-V9, and V1-V9) using V-Xtractor^76^ for training models with different region as input. Each region was converted to a 1to5-mer embedding as described above, and inherited its assembly’s EC three-digit profile.

#### Model architecture and training

embeRNA is a shallow fully connected neural network that maps a 16S rRNA k-mer embedding to EC probabilities. The architecture comprised two hidden layers (512 units each; ReLU activations) and a final linear layer producing logits for each EC label. Models were implemented in PyTorch^77^ (v2.1.2) and trained with Adam optimizer (learning rate 10⁻³; batch size 1,024) to minimize binary cross-entropy on logits (independently per label). At inference, logits were converted to probabilities using the sigmoid function.

#### Taxonomy-stratified train/validation/test splits

To evaluate generalization to unseen taxa, we performed taxonomy-stratified splits at a user selected rank (including assembly, species, genus, family, order, class, phylum). Assemblies were grouped by their lineage at that rank and randomly partitioned into train/validation/test (80/10/10); all 16S rRNA sequences of given region from assemblies in a group were restricted to the same split. Unless noted, results use genus-level stratification.

#### Model selection and binarization threshold

We trained multiple (default: 10) replicate models using different random seeds for train/validation/test split. For each replicate, the checkpoint with the lowest validation loss was selected. On each replicate’s test set, we computed precision-recall curves (on all predictions across labels and samples) and the corresponding PR-AUC.

We built the final embeRNA model by initializing from the best-performing replicate and retraining on all available 16S rRNA-function pairs in the *embeRNA set*. We used the median of threshold values where each of the ten replicate models achieved their best F1 score as the final threshold (t_m_; **SOM Table 1**) to convert embeRNA predicted values to binary (>t_m_ as presence, otherwise absence).

**Table 1.** Whole genome k-mers predict functions.

| Model | per-function F1 (mean ± SE)* | Hamming loss (mean ± SE)** |
| --- | --- | --- |
| **Logistic Regression (LR)** | **0.750 ± 0.008** | 0.136 ± 0.001 |
| **Extra Trees (ET)** | 0.714 ± 0.010 | **0.108 ± 0.002** |
| **Random-prevalence baseline** | 0.573 ± 0.011 | 0.237 ± 0.001 |
| **LR permuted-label control** | 0.562 ± 0.010 | 0.276 ± 0.001 |
| **ET permuted-label control** | 0.538 ± 0.013 | 0.183 ± 0.002 |
| **Majority baseline** | 0.511 ± 0.014 | 0.173 ± 0.001 |

Bolded values indicate best performance per column. \*Values are mean ± SE F1 (Eqn. 1) across function labels and cross-validation (CV) folds. F1 scores weigh all functions equally regardless of prevalence; \*\*Values are mean ± SE across CV folds; Hamming loss (Eqn. 2)

The entire front-to-end process of training and validating procedure, test-set evaluation, and final model selection of an embeRNA model for selected 16S rRNA region (e.g., V6-V8), at given taxonomy split (e.g., genus), can be found in **SOM Note 2**.

#### embeRNA-Fusion models

To generalize beyond EC space, we trained analogous embeRNA models to predict Fusion function from 16S rRNA 1to5-mers, using the same workflow. To reduce sparsity, the Fusion output space was restricted to functions present in ≥10% of Fusion assemblies).

### Evaluation of embeRNA on Novel Microbes set

On the *Novel Microbes set* (sequences with <97% similarity to any 16S rRNA in *embeRNA set*), we evaluated how well embeRNA can inform EC three-digit functions if the V3-V4 region of these novel 16S rRNA sequences were captured.

#### Compared 16S rRNA-based prediction strategies

Using V3-V4 as input, we compared:

- embeRNA: We applied the finalized V3-V4 genus-split model to predict functions for each sequence, binarizing the predicted values with t_m_ **(SOM Table 1)**.
- PICRUSt2: We customized the PICRUSt2 reference assembly function profiles by: (i) updating the four-digit Enzyme Commission (EC) profiles using the latest annotations from the JGI IMG database (successfully updating 19,717 of the 20,000 references); and (ii) generating a three-digit EC presence/absence profile from these updated four-digit data to serve as the PICRUSt2 reference function table. Each V3-V4 sequence was then placed into the reference tree by PICRUSt2, and functions were predicted using these customized profiles.
- Taxonomy-to-core approaches: for each sequence, we (i) first assigned taxonomy using Kraken2^39^ and RDP Classifier^59^ separately, then (ii) predicted the “core” set of EC three-digit functions present in all *embeRNA set* genomes within the assigned group; if an assigned taxon is absent from *embeRNA set*, back off to higher ranks; if no assignment/match exists, return all-zero.

#### Evaluation

Predictions across all functions and tested samples were compared to ground-truth EC profiles of *Novel Microbes set* restricted to the intersection of functional spaces supported by all tools, using precision, recall, and F1 (**Eqn. 1**).

We next compared the two best-performing tools, embeRNA and PICRUSt2, on the hard to predict functions, i.e. those for which the two approaches made opposite predictions. Using the true function presence/absence as the reference, we calculated the proportion of correct predictions for each tool across three scenarios: (i) all disagreement cases, (ii) disagreement cases in which the function was truly present, and (iii) disagreement cases in which the function was truly absent. As a baseline, we generated random presence/absence predictions by Bernoulli sampling with probability equal to the observed prevalence of truly present functions in the disagreement set (p = 0.44), and evaluated this baseline across the same three scenarios.

### Evaluation of embeRNA on real world microbiome data (*Microbiome set*)

On a set of 22 soil samples with both whole metagenome shotgun sequencing (WMS) and V6-V8 16S rRNA amplicon sequencing data, we further evaluated embeRNA’s EC three-digit function prediction against those informed from WMS by HUMAnN 3^44^.

#### Predicting functions from 16S rRNA using embeRNA

We first processed the input sequences by (i) merging paired-end reads using bbmerge^78^ (v39.06), (ii) extracting the exact V6-V8 regions using V-Xtractor (v2.0), and (iii) embedding reads to 1to5-mers. The 1to5-mers were then passed through the finalized V6-V8 embeRNA model for function prediction. Per-16S EC probabilities were binarized using t_m_ **(SOM Table 1)**. Presence for each function were then summed through all 16S sequences to obtain within sample function counts / abundance.

#### Predicting functions from 16S rRNA using PICRUSt2

We first generated amplicon sequencing variants (ASV)s and an ASV abundance table from the V6-V8 data using DADA2^18^ (v3.11). The resulting ASVs were then processed with PICRUSt2 to predict functions, using the customized EC three-digit reference function table described previously. Notably, when constructing sample-level function pseudo-abundance profiles, we omitted PICRUSt2’s default normalization by 16S rRNA gene copy number so that the resulting profiles would be directly comparable to those generated by embeRNA.

#### Predicting functions from WMS using HUMAnN 3

We quality-filtered WMS reads with fastp^79^ (v0.23.4; parameters: -e 25 -l 70) and removed host contamination by (i) aligning reads to the human reference genome (GRCh38^80^) with BWA-MEM^81^ (v0.7.17), and (ii) processing alignments with SAMtools^82^ (v1.19.2). We then used HUMAnN 3^44^ (v3.9) with the UniRef50 database^83^ to infer EC four-digit function profiles, which we summarized to EC three-digit profiles by summing abundances across all EC four-digit entries within each EC three-digit category.

#### Comparison to WMS profiles

We compared the per-sample function categories predicted by embeRNA and PICRUSt2 with those inferred from HUMAnN 3. For each 16S rRNA-based method, we quantified (i) the proportion of WMS-inferred function categories recovered in each sample and (ii) the number and proportion of additional function categories predicted by the method but not detected by HUMAnN 3.

For abundance-based comparisons, we converted each sample’s function counts to relative abundances by normalizing to the total function count in that sample. We then calculated Spearman correlations between each 16S rRNA-based function profile and its corresponding WMS-based profile.

#### Comparison of discordant function abundance bins between embeRNA and PICRUSt2

To assess differences in abundance estimation between embeRNA and PICRUSt2, we compared functions whose predicted abundance levels differed between the two tools within the same sample, and evaluated those discordant functions against the corresponding HUMAnN 3-derived abundance levels. For each sample and method (embeRNA, PICRUSt2, and HUMAnN 3), we converted function abundances to relative abundances and assigned functions to ten decile bins based on within-sample abundance rank. We then identified functions with substantial disagreement between embeRNA and PICRUSt2, defined as a difference of at least three abundance bins, and evaluated which tool assigned the abundance bin closer to that of the corresponding HUMAnN 3-derived profile. For each sample, we summarized the proportion of discordant functions for which each 16S rRNA-based tool assigned an abundance bin closer to the corresponding HUMAnN 3 bin.

### Statistical analyses

Where applicable, we estimated the 95% confidence intervals (CIs) for observed values via bootstrapping, using 80% subsamples of the data without replacement over 1000 iterations.

## Results

### Whole genome k-mer representations reflect genome-encoded functions

We first investigated whether bacterial whole genome k-mer representations (1to5-mers; **Methods**) reflect the corresponding encoded functions. Using 1,369 bacterial genomes (1,349 unique species) from the *Balanced set*, we trained classifiers to predict enzymatic function presence from whole genome k-mer vectors (**Methods**). These classifiers substantially outperformed simple (random and majority) and permuted-label control baselines (Table 1; **Methods**), confirming that genome k-mer composition carries predictive information about the encoded functional repertoires.

### 16S rRNA k-mers reflect whole genome k-mer composition

To evaluate if 16S rRNA sequences can predict whole genomes in the k-mer space, we trained regression models using the genomes in the *Balanced set* (**Methods**). Both of our models substantially outperformed the dummy mean baseline (**Table 2**) and the two permutation controls: (i) shuffled genome/16S rRNA pairings and (ii) shuffled within-genome k-mer values, confirming that predictive signal is specific to real sequence relationships.

**Table 2.** Predicting whole genome k-mers from 16S rRNA k-mers.

| *Condition* | *Model* | *R<sup>2</sup>*<br>*(mean ± SE)\** |
| --- | --- | --- |
| ***Real data*** | Ridge regression | **0.785 ± 0.008** |
|  | Random forest | **0.784 ± 0.006** |
|  | Dummy mean baseline | -0.004 ± 0.002 |
| ***Permuted pairs*** | Ridge regression | -0.007 ± 0.010 |
|  | Random forest | 0.005 ± 0.012 |
|  | Dummy mean baseline | -0.004 ± 0.002 |
| ***Genome permutation*** | Ridge regression | -9.992 ± 0.288 |
|  | Random forest | -9.834 ± 0.282 |
|  | Dummy mean baseline | -9.831 ± 0.282 |

\*Values are mean ± SE across CV folds.

### embeRNA predicts microbial functions from 16S rRNA k-mers

We further asked whether 16S rRNA-derived k-mers could directly predict genome-encoded functions.

Note that complete 16S rRNA gene sequences, comprising ten semi-conserved regions interspersed with nine hypervariable regions, are primarily inferred from complete genome sequencing. However, targeted/amplicon sequencing used for microbiome analysis relies on a collection of available primers. The longest sequence that can thus theoretically (if not practically) be extracted contains all nucleotides from just before the start of variable region 1 (V1) through the end of variable region 9 (V9), i.e. effectively dropping most of the first and last conserved regions. We thus used the 16S rRNA V1-V9, extracted from the complete genome data in our *embeRNA set* (24,585 genomes), as input and binary EC three-digit profiles as output to develop the first version of embeRNA - a neural network for predicting functions from 16S rRNA k-mer representations (**Methods**).

To rigorously evaluate model generalization, we used a genus-stratified strategy, where all genomes from the same genus were assigned to the same split (train, validation, or test). Across ten independent repetition models with different random splits, embeRNA achieved a median F1=0.91 (IQR: 0.89-0.91) on held-out test sets; best-performing replicate reached F1=0.92 (precision 0.93, recall 0.91) and precision-recall area under the curve (PR AUC) of 0.98. A shuffled-label baseline, randomly permuting labels across all samples in each test set, performed substantially worse (PR AUC = 0.62; **Methods**), confirming that embeRNA captured genuine sequence k-mer-function associations.

The final (V1-V9) model, retrained on all available data in *embeRNA set*, achieved F1=0.96 (precision 0.96, recall 0.96; **Eqn. 1**) when evaluated on the full *embeRNA set*, representing best-case performance on previously seen sequences (thresholds derived as the median optimal F1 cutoff across replicates, **SOM Table 1; Methods**).

### embeRNA generalizes across amplicon-sequenced regions

The first embeRNA model used the nearly complete 16S rRNA gene sequence (V1-V9) as input. However, most amplicon sequencing datasets focus on shorter, mostly variable, regions. V3-V4 is most frequently targeted in modern 16S rRNA amplicon sequencing studies, followed closely by the V4 region alone. V3-V4 has well-validated, nearly universal primers and spans ∼460bp, which is ideal for paired-end Illumina sequencing (e.g. 2×250bp or 2×300bp). It thus allows for high quality, cost-effective sequencing while maintaining strong taxonomic classification accuracy. The V4 region alone (∼290 bp) is commonly selected in large-scale projects for cost efficiency. Historically, V6-V8 was popular during the 454 pyrosequencing era but fell out of favor with global switch to Illumina and problematic GC-regions^84^. The choice of gene regions to sequence often depends on the sequencing platform constraints, target taxonomic groups, and plans for comparison to existing datasets.

We thus evaluated embeRNA across six additional regions (**Figure 2**). All region-specific models performed comparably to V1-V9: for example, the V6-V8 model achieved a median F1 of 0.89 [IQR: 0.88-0.90], with the best replicate reaching F1 of 0.93 and PR AUC of 0.98 (shuffled-label baseline PR AUC = 0.60). Performance was robust across all tested regions (**Figure 2**).

**Figure 2.**
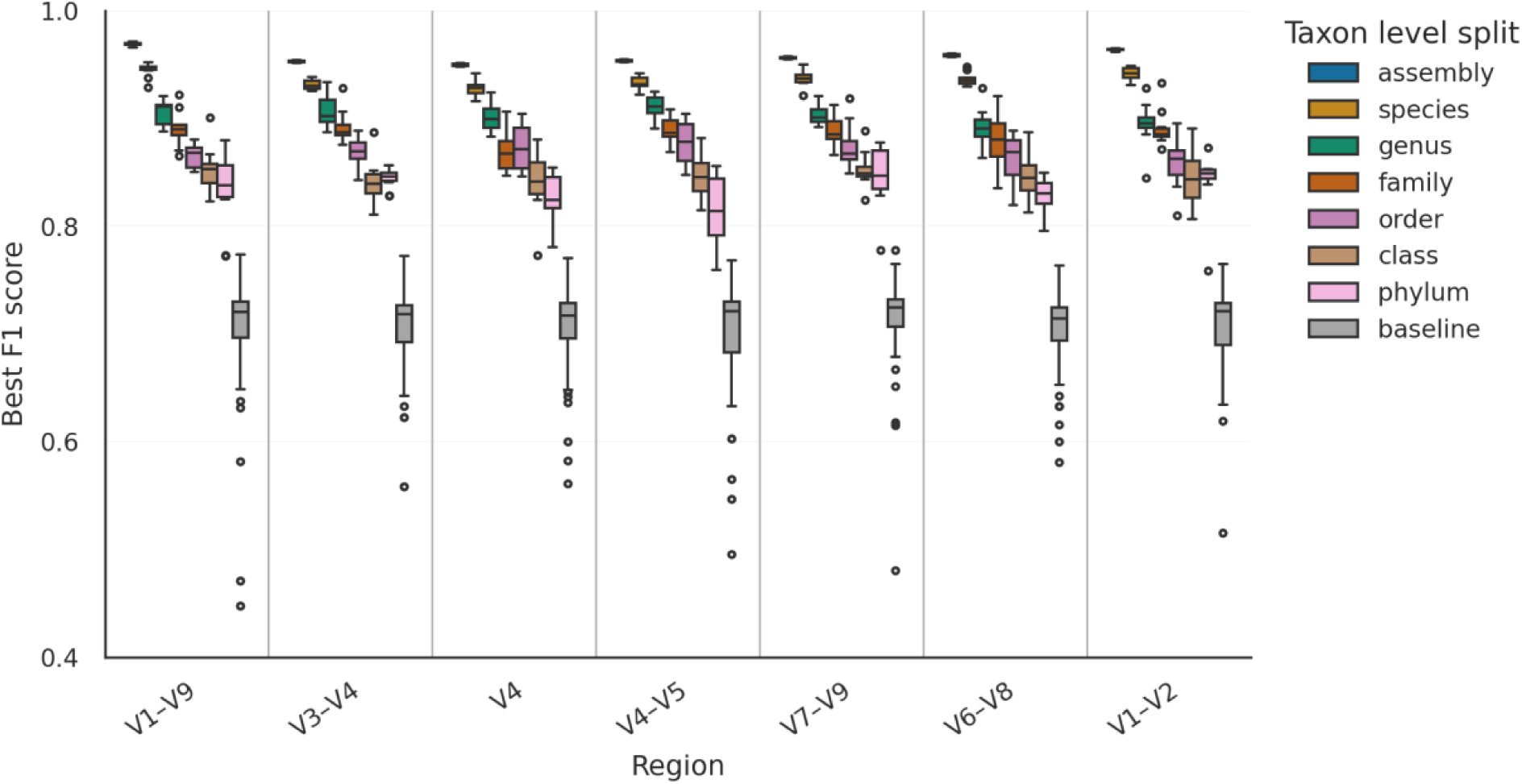
embeRNA maintains strong performance across 16S rRNA regions and stringent taxonomic holdout splits. Boxplots show the distribution of best F1 scores for embeRNA-EC models trained on different 16S rRNA regions (x-axis) and evaluated under increasingly stringent taxon-level holdout splits (colours: assembly to phylum; “baseline” indicates a random/shuffled control). For each region × split level, results are summarized across ten independent replicate models (different random seeds for splitting training/validation/testing), each evaluated on its corresponding held-out test set. Vertical separators delineate regions. Alt text: Boxplots of best embeRNA F1 scores across seven 16S rRNA regions (V1–V9, V3–V4, V4, V4–V5, V7–V9, V6–V8, V1–V2) under increasingly stringent taxonomic holdout splits from assembly to phylum level, with random baselines.

### embeRNA outperforms reference-based methods on phylogenetically novel 16S rRNAs

To evaluate functional inference on previously unseen taxa, we assembled a *Novel Microbes set* (**Methods**): 5,542 16S rRNAs from genomes released after the embeRNA training set, with <97% similarity to any training reference (median maximum similarity: 95%, IQR: 93-96%).

We asked: how well could embeRNA inform functions for these novel microbes? From the *Novel Microbes set*, V3-V4 regions were extracted (5,473 sequences) and EC three-digit functions predicted using four strategies: (i) embeRNA V3-V4; (ii) PICRUSt2 (phylogenetic placement); and (iii-iv) taxonomy-to-core approaches using Kraken2^39^ and RDP Classifier^59^ k-mer based taxonomic classifiers; taxonomy-to-core methods assign each V3-V4 sequence to a taxonomic group and use the “core” EC set of functions present in every member of the assigned group (**Methods**).

#### Prediction coverage differs across tools

By nature of the neural networks, embeRNA generates predictions for every input sequence and therefore was able to infer presence or absence of functions for all 5,473 V3-V4 sequences. For the taxonomy-to-core function approaches, both RDP Classifier and Kraken2 assigned all sequences to a taxonomic group. However, for 511 sequences (9%), the Kraken2-assigned taxa (and their ancestors) were absent from the *embeRNA set*, preventing function inference. For PICRUSt2, one sequence was too distant from the reference tree to yield predictions, and four additional sequences had NSTI > 2 (nearest sequenced taxon index), a range suggested to generate unreliable predictions due to lack of closely related references^40^; otherwise, median NSTI=0.06 [IQR: 0.03-0.11].

For standardizing the comparison data, analysis was restricted to the 4,961 V3-V4 sequences where all four tools generated predictions. The downstream evaluation functional space was defined as the union of 255 functions spanning (i) the 237 functions predictable by all four 16S rRNA-based tools and (ii) the 241 actual functions across *Novel Microbes set* assemblies. The microbes in this set contain a median of 137 unique EC three-digit functions each. embeRNA and PICRUSt2 predicted comparable numbers of functions per sample - a median of 137 and 140, respectively, versus 86-89 for taxonomy-to-core approaches.

#### embeRNA attained best overall performance

embeRNA achieved the highest F1 of 0.851 (95% bootstrap CI: [0.851-0.852]), followed by PICRUSt2 (0.835; CI: [0.834-0.835]), Kraken2 (0.751; CI: [0.749-0.753]), and RDP (0.749; CI: [0.748-0.751]). The taxonomy-to-core methods achieved highest precision (∼0.93) but substantially lower recall (∼0.63), consistent with their conservative “core functions only” design. embeRNA and PICRUSt2 attained relatively lower precision (embeRNA precision=0.845; PICRUSt2 precision=0.816), and higher recall (embeRNA recall=0.857 and PICRUSt2 recall=0.854).

#### embeRNA enables tunable precision-recall trade-offs

Unlike the binary outputs of other tools, embeRNA outputs continuous probabilities (**Figure 3**), enabling threshold adjustment to prioritize precision or recall. For example, at RDP-precision-matched operating point (precision = 0.927), embeRNA achieved substantially higher recall (0.716, 95% CI: [0.713-0.720]) vs. RDP (0.629).

**Figure 3.**
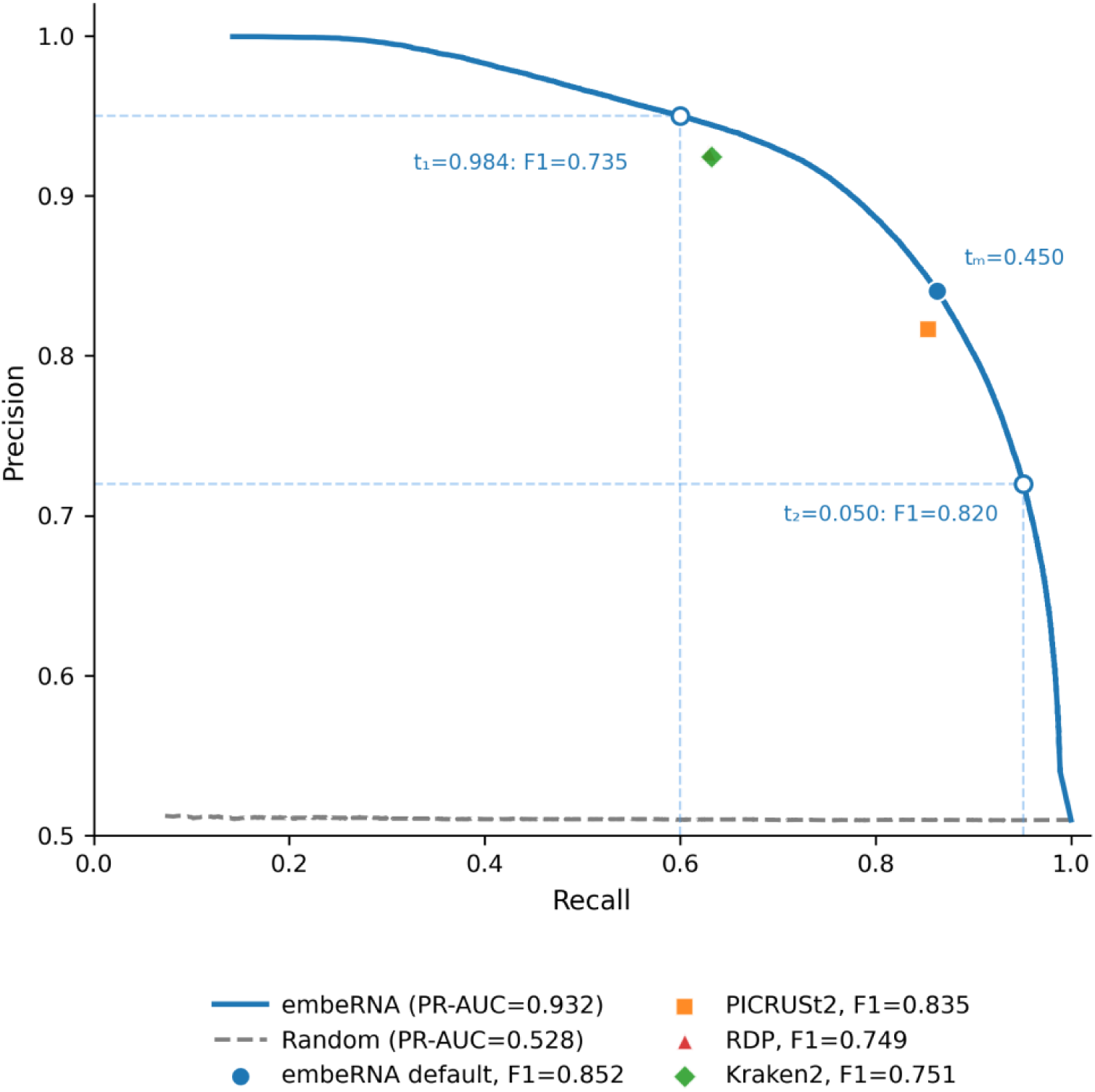
embeRNA outperforms reference-dependent 16S rRNA-based approaches on the *Novel Microbes set* and permits threshold-dependent tuning. Precision-recall (PR) curve for embeRNA-EC V3-V4 model evaluated on the *Novel Microbes set* (PR-AUC = 0.932). The grey dashed line shows the random shuffled baseline (PR-AUC = 0.528). Filled markers denote the default operating points for embeRNA (t_m_=0.450: F1=0.852, precision=0.841, recall=0.863), PICRUSt2 (F1=0.835), RDP taxonomy-to-core (F1=0.749), and Kraken2 taxonomy-to-core (F1=0.751) on the same V3-V4 sequences. Open blue circles indicate two additional embeRNA operating points obtained by adjusting the classification threshold to favour either higher precision (with threshold t_1=_0.984: F1=0.735, precision=0.950, recall=0.600) or higher recall (with threshold t_2_=0.050: F1=0.820, precision=0.720, recall=0.951); dashed guidelines project these points to the axes to illustrate the associated precision-recall trade-offs. Alt text: Precision–recall curve for the embeRNA V3–V4 model on the *Novel Microbes set*, with the embeRNA default operating point and fixed-point comparisons for PICRUSt2, RDP and Kraken2 taxonomy-to-core methods. Two additional threshold-tuned operating points illustrate precision–recall trade-offs for embeRNA.

#### For hard to classify functions embeRNA has fewer false positives than PICRUSt2

Some functions such as transcription and translation are present everywhere and therefore easy to predict. Others are absent across the vast swaths of the community and are also fairly easy to label as such. We asked about the difficult ones, i.e. when embeRNA and PICRUSt2 give conflicting calls for the same function, which is more likely to be correct?

As taxonomy-to-core approaches called less than two thirds of the actual functions, we restricted this analysis to embeRNA vs PICRUSt2 to avoid conflating disagreement with coverage differences. Here, we used each genome’s encoded functions as ground truth for each prediction.

Among the 27 (median, [IQR: 21-35]) functions per 16S rRNA, where embeRNA and PICRUSt2 disagreed (159,618 total disagreement calls across all *Novel Microbes set*), embeRNA was more frequently correct than PICRUSt2 (58.6% of the time vs. 41.4%). Note, a random baseline was correct 50.7% of the time (**Methods**). This advantage was most pronounced for truly absent functions (true negatives, n = 89,547; embeRNA correct 63.9% of the time vs. PICRUSt2 36.1%; random baseline 56.0%), indicating fewer false-positive calls in the hard to predict functional space (**Figure 4**, **SOM Figure 2**). Among truly present functions (true positives, n = 70,071), embeRNA remained more accurate than PICRUSt2 (52.0% vs. 48.0%; 43.8% for random), although slightly less obviously so. This true-negative advantage is orthogonal to F1, which by definition does not capture absence-calling (negative) accuracy.

**Figure 4.**
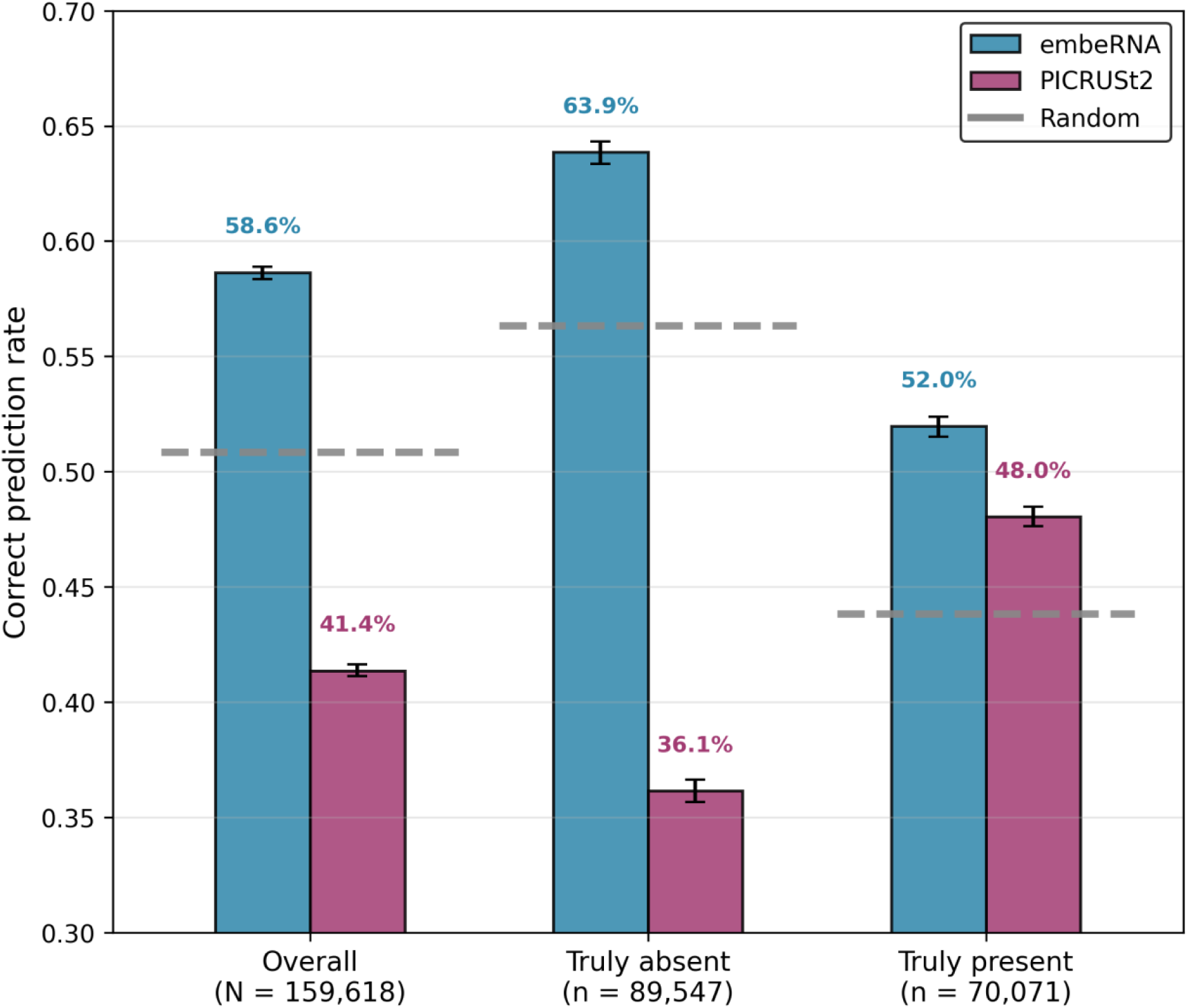
embeRNA outperforms PICRUSt2 in head-to-head prediction of hard to predict functions in the *Novel Microbes set*. Bar plots show the proportion of correct predictions made by each tool (embeRNA, blue; PICRUSt2, purple) among hard to predict functions for which the two tools disagreed on presence or absence. Error bars indicate 95% bootstrap confidence intervals. The random baseline (grey dashed lines) was generated by assigning function presence or absence according to each function’s empirical prevalence. Alt text: Bar plot comparing correct prediction rates of embeRNA and PICRUSt2 for hard-to-predict functions where the two tools disagreed on *Novel Microbes set*.

### embeRNA predicts WMS-inferred functions in soil samples well

We compared functional profiles from embeRNA and PICRUSt2 against HUMAnN 3^44^ processed WMS data for 22 blueberry rhizosphere and bulk soil samples with paired 16S rRNA V6-V8 amplicon and WMS sequencing data^75^ (**Methods**). Both 16S rRNA-based tools recovered 95% of WMS-detected functions per sample, as expected given the limited depth of WMS vs. amplicon sequencing^57^.

Additionally, embeRNA and PICRUSt2 predicted a quarter more functions than WMS (median embeRNA 234, PICRUSt2 235, versus HUMAnN 3 186 EC functions per sample). Most (56 per sample) of these functions were predicted by both 16S tools.

Note that in microbiome studies, abundance profiles are more common than presence/absence profiles. We computed pseudo-abundance profiles by scaling embeRNA’s and PICRUSt2’s per-16S presence/absence vectors according to their observed frequencies within a sample (**Methods**). embeRNA’s abundance profiles were better correlated with HUMAnN 3 (Spearman correlation median ρ = 0.74, IQR: 0.72-0.75) than PICRUSt2’s profiles (median ρ = 0.70, IQR: 0.69-0.71).

To understand performance differences, we further compared the tools’ function abundance levels by (i) determining each function’s relative abundance within each tool’s profile, and (ii) splitting functions in each sample into ten decile bins based on their abundance rank. Across the compared functions per sample (median 251), for functions with large abundance bin discrepancies between the two tools (≥3 bins apart; ∼4% of compared functions per sample), embeRNA’s estimates aligned more closely with WMS in a median of 72% of cases per sample (IQR: 64-82%; for example, sample Bact200 is displayed in **Figure 5**).

**Figure 5.**
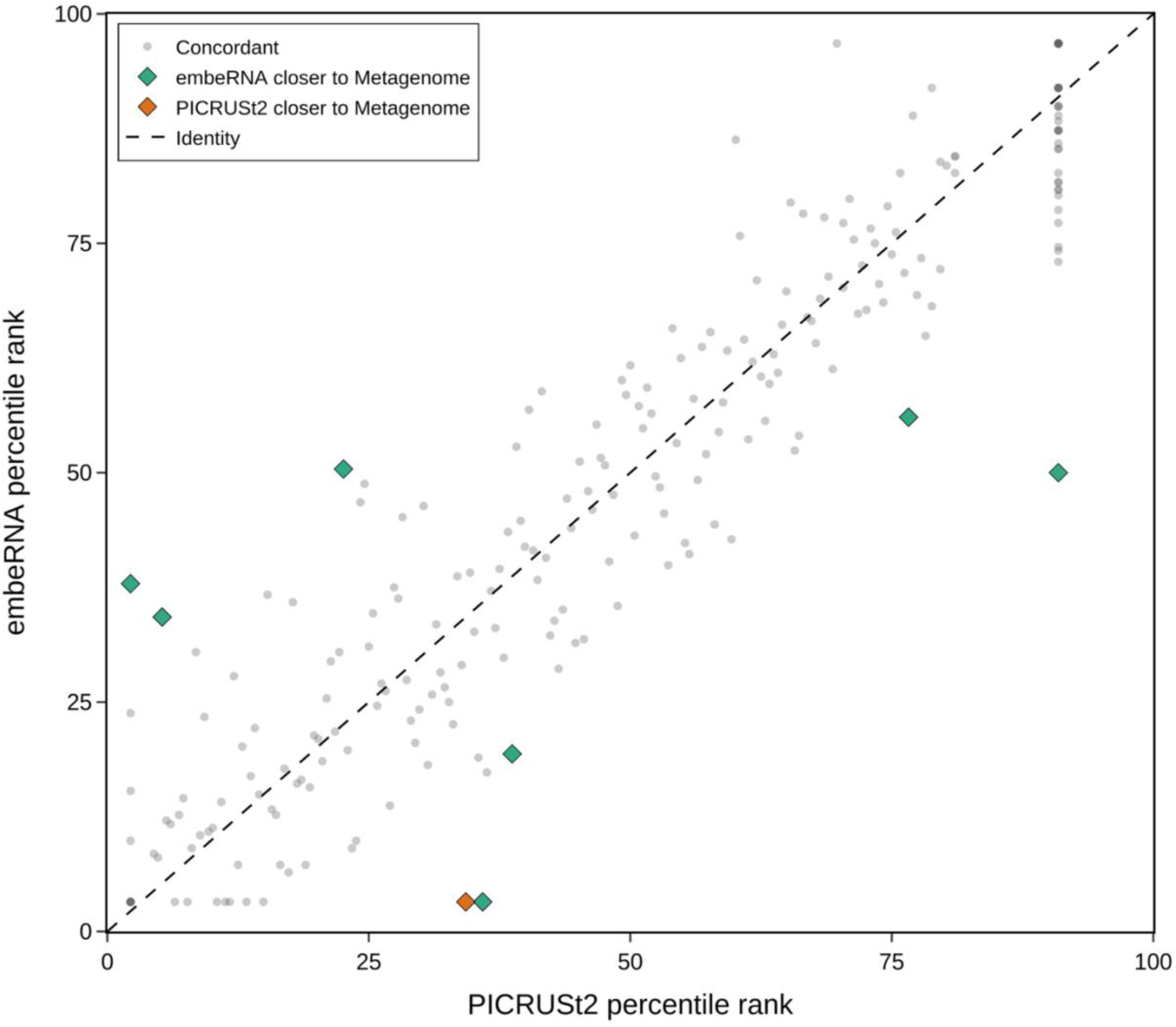
embeRNA more accurately reflects WMS-informed functional abundance when predictions disagree with PICRUSt2. Shown is an example soil sample (Bact200) with paired 16S rRNA amplicon and whole-metagenome shotgun (WMS) data. Functional profiles were inferred from 16S data using embeRNA and PICRUSt2, and from WMS using HUMAnN3. Within each method, functions were ranked and grouped into decile-based abundance bins. Each point represents a function, positioned by its percentile rank in the PICRUSt2 profile (x-axis) and embeRNA profile (y-axis). The dashed line indicates identity between methods. Functions with substantial disagreement (≥3-bin difference) were evaluated against the WMS-derived abundance: points are colored green if embeRNA is closer to WMS, orange if PICRUSt2 is closer, and blue if both are equally close. Alt text: Scatter plot comparing functional abundance percentile ranks from embeRNA (y-axis) and PICRUSt2 (x-axis) for a soil sample. Most functions cluster near the identity line, indicating agreement. Among functions with large disagreements, green diamonds (embeRNA closer to WMS) substantially outnumber orange diamonds (PICRUSt2 closer), with only one orange point visible.

### embeRNA framework is flexible with respect to the choice of functional space

Although until this point we focused on EC three-digit functions, embeRNA can be trained to predict any user-defined functions. The Fusion database, for example, catalogs 433,891 microbial functions, spanning both known and uncharacterized functionality^71^. We trained embeRNA-Fusion models using 16S rRNA sequences from the Fusion-specific assemblies paired with curated Fusion functional profiles. To reduce sparsity and focus on broadly supported functions, we trained embeRNA with, and predicted, only the Fusion functions present in at least 10% of Fusion assemblies (4,482 functions total).

Across ten repeated random genus-level split evaluations using full-length (V1-V9) 16S rRNA, embeRNA-Fusion achieved a median F1 of 0.82 on held-out test sets (precision 0.81; recall 0.84; **SOM Figure 3**). Models trained on shorter regions performed comparably (e.g. V6-V8 F1 0.80; precision 0.80; recall 0.81), demonstrating that the same 16S rRNA k-mer-to-function framework readily adapts to alternative ontologies and customized function sets.

## Discussion

In this work, we describe a link between 16S rRNA and organismal function that has not previously been observed: 16S rRNA k-mer composition reflects that of its source genome, which, in turn, tracks the history of and organism’s evolutionary and environmental constraints - the two factors that also shape its function potential. Based on this observation, we built embeRNA, a framework that infers functional potential from 16S rRNA k-mer composition modelling rather than from taxonomy or phylogenetic placement, enabling functional characterization for microbial “dark matter”.

### Mechanistic basis for inferring function from 16S rRNA k-mer composition

A central question underlying embeRNA is whether (and why) the k-mer composition of a single marker gene carries sufficient information to infer an organism’s broader functional potential. Our results support a mechanistic bridge built on two principles of genomic evolution.

First, whole genome k-mer composition is predictive of genome-encoded functional profiles. This relationship is governed by two interacting forces: ancestral lineage and environmental selection. Lineage history provides the primary template for genomic signatures, as inherited biases in replication and DNA repair mechanisms drive characteristic oligonucleotide usage patterns^66–69^. These ‘phylogenetic signals’ ensure functional conservation within lineages^85, 86^. However, prolonged exposure to specific ecological constraints further refines these patterns, driving functional convergence across disparate lineages^87, 88^. Habitat-linked shifts in GC content and higher-order oligonucleotide properties demonstrate how environmental filtering imposes readable, niche-specific signatures on the genome^69, 89–91^. Because lineage history and niche adaptation simultaneously shape functional repertoire and genome k-mer composition, the latter serves as a robust proxy for the former.

Second, 16S rRNA k-mer composition closely mirrors that of its source genome, consistent with the established cohesion of compositional signatures across genomic regions^66–68, 90^. Together, these two observations close a logical chain: genome k-mers predict functions, and 16S rRNA k-mers reflect genome k-mers, so direct prediction from 16S rRNA k-mers to functions is mechanistically grounded. This chain supports generalisation beyond taxonomic boundaries: even when a query sequence has no close phylogenetic neighbour in the training data, the signal embedded in its k-mer composition shaped by lineage and environmental adaptation history may still point toward the correct functional archetype.

### embeRNA’s compositional inference reduces false positives for distant relatives

embeRNA’s small advantage over PICRUSt2 on novel microbes (F1: 0.851 vs. 0.835) understates the most consequential method difference, which lies in the true-negative calls that the F1 metric fails to capture. Among functions where the two tools disagree, embeRNA correctly identifies truly absent functions twice as often as PICRUSt2; that is embeRNA says these are absent, while PICRUSt2 claims their presence.

This pattern extends to community-level analysis: in soil samples, among functions where embeRNA and PICRUSt2 disagree, embeRNA’s abundance estimates aligned more closely with WMS 72% (median, IQR: 64%-82%) of the time. The convergence of this pattern across the set of individual novel microbes and complex soil communities supports the robustness of embeRNA’s compositional inference.

The mechanistic basis for this reduced false-positive rate lies in how the two tools handle novelty. PICRUSt2 transfers functional annotations from the nearest reference(s) in a pre-built phylogenetic tree. For divergent sequences, the nearest reference may be taxonomically distant, and imputation produces dense predictions reflecting the reference’s broad functional repertoire rather than the query organism’s specific capabilities, systematically overcalling reference-specific traits. embeRNA, by contrast, assigns per-function probability scores anchored to the query’s own k-mer signature: when a functional signal is absent from the k-mer profile, the model assigns low support regardless of what phylogenetic neighbours encode.

A practical consequence is the tunability of embeRNA’s output. Because predictions are continuous probabilities rather than binary calls, users can adjust the decision threshold to match study-specific precision-recall requirements. At precision-matched operating points (precision = 0.927, matching RDP), embeRNA achieved higher recall (0.716 vs. 0.629), demonstrating genuine flexibility rather than a fixed trade-off.

### Complementarity between WMS and 16S rRNA-based function inference

Both embeRNA and PICRUSt2 recovered WMS-inferred soil functional profiles well, but both also predicted additional functions per sample (a median of 58 for embeRNA and 64 for PICRUSt2). Some of these are ecologically central to soil biogeochemical cycles: nitrogenases (EC 1.18.6 and EC 1.19.1, predicted by both tools in all samples) and complex carbohydrate degraders (EC 4.2.2, predicted by both tools in 13 samples) are well-documented components of soil microbial communities^92, 93^. We thus suspect that WMS sequencing depth may have excluded these likely true positives^57^. However, as embeRNA and PICRUSt2 share overlapping reference genome collections, we cannot exclude the chance that these agreements reflect shared systematic over-predictions. Resolving the true source of these observations would require deeper WMS sequencing or targeted experimental validation.

Whereas embeRNA and PICRUSt2 are expected to perform similarly well for well-characterized organisms, our results suggest that embeRNA provides a greater advantage for predicting functions in novel microbes with limited reference coverage.

### Expanding functional inference into unannotated space

The embeRNA architecture is inherently flexible: to illustrate this, we trained embeRNA-Fusion on the Fusion database and applied it to the same 22 blueberry soil samples^75^, for which no Fusion-based profiling had previously been reported.

Predicted functional profiles clustered by sampling site (PERMANOVA: R²=0.42; p=0.0003) and, within the Collingwood site, by soil compartment (rhizosphere vs. bulk; R²=0.33, p=0.01; **SOM Note 3**). Differentially abundant functions between rhizosphere and bulk soil (FDR < 0.05) included enrichment of functions describing chemical defence and physical colonization machinery in the rhizosphere which involve functions not described in the Enzyme Commission space, validating the framework against known root-interface biology^94–97^.

More notably, 43% of the differentially abundant functions corresponded to Fusion clusters with no annotated protein sequences, demonstrating that embeRNA-Fusion detects ecologically structured variation in functionally dark space, a capability that EC-based approaches cannot provide. Experimental validation of these uncharacterised functions, for example, via metatranscriptomics or targeted biochemical assays, represents an important direction for future work. We expect embeRNA to provide novel understanding of microbial functions in other underrepresented ecological microbiomes.

## Conclusions

We demonstrate that 16S rRNA k-mer composition preserves genomic signatures predictive of bacterial genome-encoded functional potential. By leveraging this link, embeRNA achieves robust performance on phylogenetically novel organisms, with a particular advantage in reducing false-positive functional assignments compared to reference-based approaches. Our results reveal complementarity between 16S rRNA-and WMS-based functional profiling: the two approaches recover largely overlapping functional landscapes, but 16S-based inference extends visibility into rare and low-abundance taxa that WMS routinely under-samples. The generalisation to uncharacterised functional space via Fusion further enables detection of ecologically structured variation in regions inaccessible to annotation-dependent approaches.

While embeRNA treats individual 16S rRNA sequences as independent inputs, microbial ecosystems are defined by emergent community-level activities. Integrating all 16S rRNA sequences from a sample into a unified input to model community-scale metabolic networks represents a natural next step. By shifting from taxonomic “guilt-by-association” to intrinsic sequence-to-function mapping, embeRNA provides a scalable framework for deciphering the functional architecture of complex microbiomes.

## Data availability

Assembly accessions for the *Balanced set*, assembly accessions and 16S rRNA identifiers for the *Novel Microbes set*, EC abundance percentile rank plots between embeRNA and PICRUSt2 for all soil samples, together with the lists of embeRNA predicted Fusion functions significantly different between soil compartments (rhizosphere vs. bulk), are available on Figshare (DOI: 10.6084/m9.figshare.31297531). Blueberry soil microbiome sequencing data are available from the NCBI Sequence Read Archive under BioProjects PRJNA434066 (16S rRNA amplicon) and PRJNA484230 (whole-metagenome shotgun)^75^.

## Software availability

The embeRNA source code is available on Bitbucket at https://bitbucket.org/bromberglab/emberna.

## Supplementary data statement

Supplementary Data are available at *NAR* Online.

## Acknowledgements

We are grateful to R. Prabakaran for all helpful discussions and to Sergio Gramacho and Aiden Maloney-Bertelli for technical support (all Emory University). We also thank all those who deposit their experimental results into public databases and those who maintain the databases.

## Funding

This work was supported by the National Institute of Dental and Craniofacial Research [R01-DE032216], and the National Science Foundation award [#2310114].

## Author Contributions Statement

Jia Liu (Data curation [lead], Formal analysis [lead], Investigation [lead], Methodology [lead], Validation [lead], Visualization [lead], Writing – original draft [lead], Writing – review & editing [equal]); M. Clara DePaolis Kaluza (Data curation [support], Investigation [support], Methodology [equal], Writing – review & editing [support]); Yana Bromberg (Conceptualization [lead], Data curation [equal], Funding acquisition [lead], Methodology [equal], Validation [equal], Supervision (lead), Writing – review & editing [lead]).

## Notes

https://doi.org/10.6084/m9.figshare.31297531

### Competing Interest Statement

The authors have declared no competing interest.

### Summary of Updates

Slightly changed the title; Corrected one of the author's names; Shortened abstract; Changed the orders of Materials and Methods section and Results sections.

