## Supplemental Material for "16S rRNA sequence captures microbial functional potential"

### Supporting online material
### for:
### 16S rRNA captures microbial functional potential

**Jia Liu & Yana Bromberg**

**Table of Contents for Supporting Online Material:**

**Supplementary Figures (S1-S3)**

**Supplementary Tables (S1)**

**Supplementary Notes (S1-S3)**

**References for Supporting Online Material**

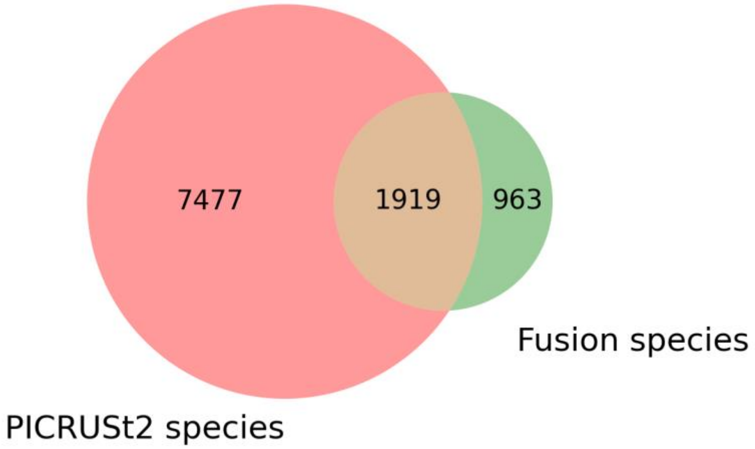


**(A)**

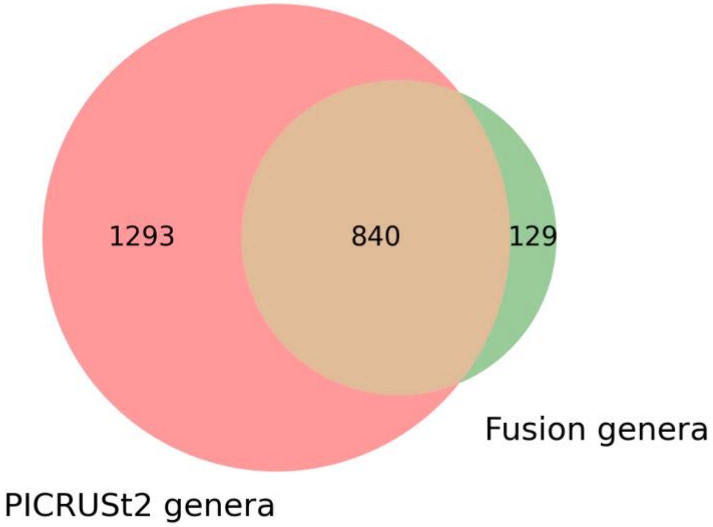


**(B)**

**Fig. S1. Taxonomy coverage of genome assemblies used to train embeRNA.**
embeRNA was trained on 24,585 prokaryotic genome assemblies**,** comprising 16,136 assemblies from the PICRUSt2 reference collection and 8,449 assemblies from the Fusion database. Venn diagrams summarize the overlap and database-specific coverage of unique taxonomic labels at the species level (A) and genus level (B).

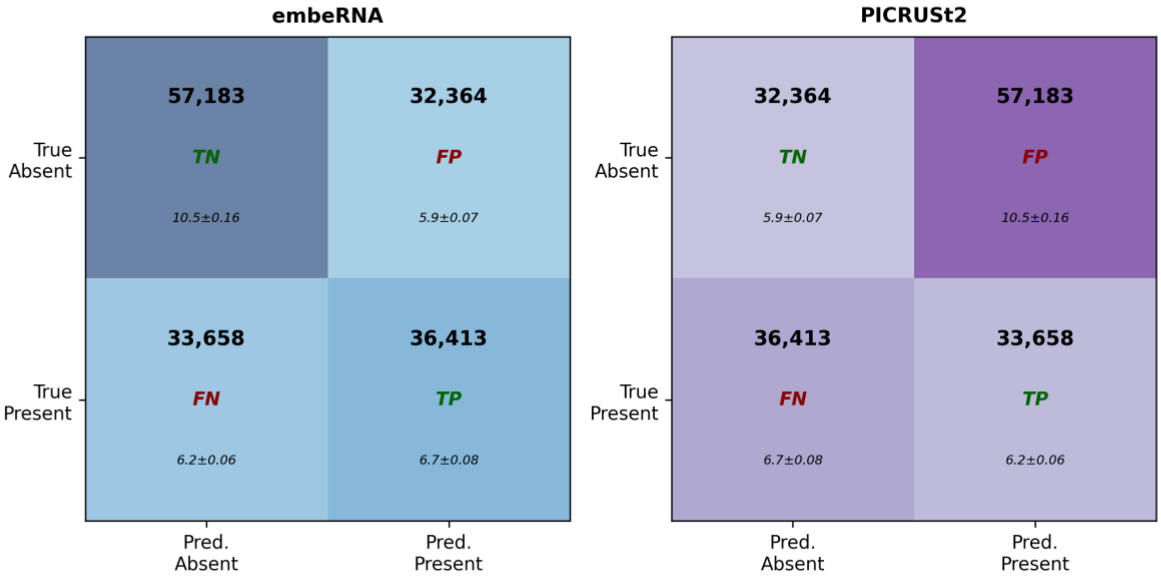


**Fig. S2. Confusion matrices across disagreed functions of *Novel Microbes set* between embeRNA and PICRUSt2.**
Classification performance of embeRNA and PICRUSt2 on the n = 159,618 hard to predict functions on which the two methods disagree of their presence/absence, evaluated against genome-derived ground truth. Each cell shows: total prediction count (top, bold), classification category (middle; TN: true negative, FP: false positive, FN: false negative, TP: true positive), and per-sample mean ± SE (bottom, italic).

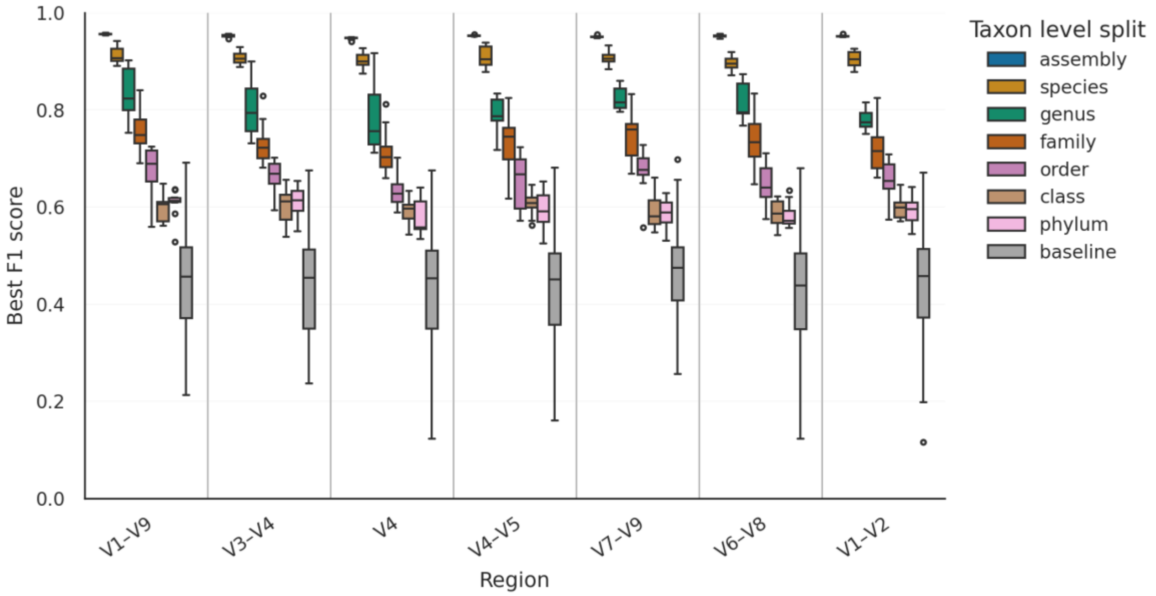


**Fig. S3. embeRNA-Fusion models maintain strong performance across 16S rRNA regions and stringent taxonomic holdout splits.**
Boxplots show the distribution of best F1 scores for embeRNA-Fusion models trained on different 16S rRNA regions (x-axis) and evaluated under increasingly stringent taxon-level holdout splits (colours: assembly to phylum; “baseline” indicates a random/shuffled control). For each region × split level, results are summarized across ten independent replicate models (different random seeds for splitting training/validation/testing), each evaluated on its corresponding held-out test set. Vertical separators delineate regions.

**Table S1. Presence/absence thresholds for embeRNA predictions across 16S rRNA regions and function spaces.**

| ***Input 16S rRNA region*** | ***embeRNA-EC**** | ***embeRNA-Fusion**** |
|---|---|---|
| ***V1-V2*** | 0.503 | 0.388 |
| ***V1-V9*** | 0.494 | 0.388 |
| ***V3-V4*** | 0.450 | 0.336 |
| ***V4*** | 0.407 | 0.326 |
| ***V4-V5*** | 0.428 | 0.384 |
| ***V6-V8*** | 0.472 | 0.357 |
| ***V7-V9*** | 0.512 | 0.295 |

*Values are medians across replicate models (probability cutoffs used to convert continuous predictions to binary presence/absence).

#### Supplementary Note 1. Construction of Novel Microbes set

We first retrieved 35,556 complete RefSeq bacterial assemblies (201,649 16S rRNA sequences) that contained at least one 16S rRNA gene and were released after February 28, 2018, i.e. a date after both components of the embeRNA set – Fusion and PICRUSt2 databases – were complete. After removing exact duplicates using seqkit{Shen, 2024 #30} (v2.7.0), 50,055 unique 16S rRNAs were retained.

To determine which of these newly released 16S rRNAs had no close relatives in the embeRNA training set, we combined these 50,055 unique sequences with all 16S rRNAs full hyper-variable region (V1-V9) sequences in embeRNA set and performed a structure-aware multiple sequence alignment (MSA) using ssu-align{Nawrocki, 2009 #29} (v0.1.1), followed by masking (ssu-mask{Nawrocki, 2009 #29}, -pf 0.001, -pt 0) to remove low-confidence positions. From the MSA output, for each 16S rRNA from newly released assemblies, we computed pairwise sequence identity to each of the 16S rRNA sequences in the embeRNA set, excluding terminal regions at both ends where either sequence in the pair contained continued gaps. Using the widely adopted 16S rRNA species-level threshold (97% identity), we identified 5,542 sequences whose maximum identity to any embeRNA set 16S rRNAs (including both Fusion and PICRUSt2) was <97%, designating them as phylogenetically novel to embeRNA set and PICRUSt2 references. These 5,542 sequences originated from 2,533 unique assemblies.

**Supplementary Note 2. Training, validation, testing, model finalizing of embeRNA V6-V8 model at genus-level split.**

Training procedure, model selection, and test-set evaluation. We trained multiple replicate models (*m*; by default: ten) using different random seeds for splitting training, validation, and test sets. Each model was trained for an initial of 500 epochs using mini-batch Adam (learning rate $10^{-3}$, batch size 1,024) optimization. After each epoch, we computed the binary cross-entropy loss on the validation set. We tracked the minimum validation loss observed and stored the corresponding model parameters and epoch index.

After the initial 500-epoch phase, we entered a fine-tuning phase in which training continued on the same train/validation split. The fine-tuning phase was (i) ran for at least N/2 additional epochs, where N is the epoch index of the minimum validation loss stored previously and (ii) allowed further training until no improvement in validation loss was observed for a fixed number of consecutive epochs (fine-tuning patience). In practice, the fine-tuning loop continued while either the minimum required number of additional epochs had not yet been completed or the number of consecutive non-improving epochs remained below the patience threshold (default 25). Whenever a new minimum validation loss was observed, we updated the stored best model parameters and reset the patience counter.

At the end of the training and fine-tuning phases, we restored the model parameters corresponding to the epoch with the lowest validation loss observed over the entire run. This model was taken as the final model for that split and seed. We would have *m* such models.

Test-set evaluation and threshold selection. Each of the *m* models was then evaluated once on its held-out test set. We obtained sigmoid probabilities for each function label and sample. We computed precision-recall curves for predictions across all labels and all test samples by comparing them to the actual function labels. From these curves we calculated the precision-recall area under the curve (PR AUC). For each model's curve, we also identified the threshold that maximized the F1-score on the test set and stored it together with (i) the best F1-score and (ii) recall and (iii) precision at that threshold. The median of this threshold across the *m* models ($t_{median}$) was reported as the reference threshold of this model's predictions.

*Finalizing the embeRNA model at given region and taxonomy split.* Based on the *m* models' performances on their test sets, we chose the one with the highest F1, and further fine-tuned it on all data in the embeRNA set, i.e. using all data as a single training set without holding out for validation or testing, for an additional 100 epochs, using the same optimizer and learning rate. The resulting model is the final embeRNA model for the given 16S rRNA region (e.g. V6-V8).

**Supplementary Note 3. embeRNA-Fusion analysis in blueberry soils.**

We applied the finalized V6-V8 embeRNA-Fusion model to infer Fusion functional profiles from soil 16S rRNA sequences. Per-sample Fusion profiles were obtained by summing per-sequence binary calls into counts and normalizing to relative abundances.

*Community-level comparisons*. Bray-Curtis dissimilarities were computed from relative abundances and tested using PERMANOVA (9,999 permutations) for site differences across all samples and, within Collingwood, for compartment, management, and their interaction (tested sequentially).

*Differential abundance*. Within Collingwood, function-wise differences between rhizosphere and bulk soils were tested using linear models on centered log ratio (CLR)-transformed relative abundances with a small pseudocount added prior to log transform. Models included compartment as the primary predictor and management as a fixed-effect covariate. P-values for the compartment coefficient were corrected across functions using Benjamini-Hochberg FDR ($FDR<0.05$). Effect sizes were on the CLR scale and converted to fold-change relative to each sample's geometric-mean functional baseline as $\exp(\beta)$.
